# On the feasibility of mining CD8+ T-cell receptor patterns underlying immunogenic peptide recognition

**DOI:** 10.1101/118539

**Authors:** Nicolas De Neuter, Wout Bittremieux, Charlie Beirnaert, Bart Cuypers, Aida Mrzic, Pieter Moris, Arvid Suls, Viggo Van Tendeloo, Benson Ogunjimi, Kris Laukens, Pieter Meysman

## Abstract

Current T-cell epitope prediction tools are a valuable resource in designing targeted immunogenicity experiments. They typically focus on, and are able to, accurately predict peptide binding and presentation by major histocompatibility complex (MHC) molecules on the surface of antigen-presenting cells. However, recognition of the peptide-MHC complex by a T-cell receptor is often not included in these tools. We developed a classification approach based on random forest classifiers to predict recognition of a peptide by a T-cell and discover patterns that contribute to recognition. We considered two approaches to solve this problem: (1) distinguishing between two sets of T-cell receptors that each bind to a known peptide and (2) retrieving T-cell receptors that bind to a given peptide from a large pool of T-cell receptors. Evaluation of the models on two HIV-1, B*08-restricted epitopes reveals good performance and hints towards structural CDR3 features that can determine peptide immunogenicity. These results are of particularly importance as they show that prediction of T-cell epitope and T-cell epitope recognition based on sequence data is a feasible approach. In addition, the validity of our models not only serves as a proof of concept for the prediction of immunogenic T-cell epitopes but also paves the way for more general and high performing models.

## Introduction

Immunoinformatics strives to computationally explore the increasingly large amounts of available immunological data by providing researchers with the necessary tools to gain novel insights into key processes of the immune system. The necessity of such immunoinformatics tools becomes particularly apparent in light of the huge complexity that underlies essential immunological processes. As the immune system has to be able to recognize a vast repertoire of non-self epitopes, it has adopted several strategies to cope with the wide range of pathogens and pathogen-derived epitopes it might come into contact with. To mount an adequate defence, the activation of the adaptive immune system requires recognition of these pathogen-derived epitopes by TCRs. Epitopes from within the cell and the extracellular environment are respectively bound by MHC class I and MHC class II molecules (Jensen 2007). The peptide-MHC (pMHC) complex is subsequently presented on the surface of the cell, where it can be recognized by the TCR of circulating CD8+ T-cells (for MHC-I) or CD4+ T-cells (for MHC-II) (Rossjohn et al. 2015). A cascade of downstream immunological pathways will then be triggered within the T-cell with the goal of eliminating the invading pathogen from which the epitope was derived. TCRs are able to bind such a wide variety of pMHC complexes due to the genetic recombination of the V and J regions in the TCR’s *α* chain and the V, D and J regions in the TCR’s β chain (Krangel 2009). These recombination events results in an estimated 10^15^ possible different TCRs (Turner et al. 2006).

Both antigen processing and presentation by MHC molecules are well-studied processes and have been documented in detail for both MHC class I and MHC class II molecules. A range of immunoinformatics tools have addressed the fundamental question of which peptides will be presented by a certain MHC molecule (Soria-Guerra et al. 2015). Several of these tools are able to predict putative epitopes with high accuracy. Furthermore, they often account for biologically relevant pre-processing steps such as proteasomal cleavage of proteins and transport of peptides into the endoplasmic reticulum by TAP transporters (Stranzl et al. 2010). Despite the diversity of possible pMHC combinations, these tools offer researchers a reliable way of setting up focused immunogenicity experiments by reducing the number of peptides that need to be experimentally tested (7-11). The success of these prediction tools stems from both our intimate understanding of the underlying biochemical processes as well as from the large amounts of pMHC affinity data that are available in public repositories such as the Immune Epitope Database (www.iedb.org) (Vita et al. 2015). However, it is important to note that while it is required that immunogenic peptides are presented to T-cells by an MHC molecule, this is not sufficient to warrant recognition by a TCR and subsequently elicit an immune response. Although these pMHC prediction methods claim to predict T-cell epitopes, they do so without any knowledge or contribution from the T-cell repertoire. These prediction tools are mainly able to differentiate between MHC-bound peptides, which could potentially be recognized by a TCR, and those peptides that are not bound by the specific MHC molecule under investigation. However, no such prediction tools exist for TCR-sequences and a given MHC-bound peptide and it is a concern if such predictions are even possible given the complexity of the recognition and the lower quantity of data.

Previous research has demonstrated that there is a differential contribution of the amino acid position in the epitope to its immunogenicity (Calis et al. 2013). As the CDR3 region of the TCR is known to interact with the MHC presented peptide (Jorgensen et al. 1992), it is to be expected that structural determinants within this region also contribute significantly to peptide recognition. In this study, we demonstrate the feasibility of constructing accurate TCR-epitope recognition predictors based on the amino acid sequence of the TCR protein. We explore the patterns underlying the interaction between peptides and TCRs, focusing on those patterns within the CDR3 region that determine epitope recognition.

## Results

Data on peptide-TCR interactions was collected from Costa et al. (2015)for two well-defined and dominant HLA-B*08-restricted HIV-1 epitopes. Control data, consisting of CD8+, HLA-B*08-restricted TCRβ sequences, was retrieved from the ImmuneACCESS database. For each dataset, the following descriptive statistics were calculated: total number of TCRβ sequences, unique CDR3 sequences, V/J families and V/J genes and the Shannon-Wiener diversity of CDR3 sequences, V/J families and V/J genes (table 1). Higher Shannon-Wiener diversity values reflect a more uniform population and/or a population with more unique samples. CDR3 sequences are the most diverse component in all datasets, followed by V gene diversity, J gene diversity, V family diversity and finally J family diversity. While the majority of CDR3 sequences that occur in a given dataset are unique, some CDR3 sequences do frequently reoccur, though the number of reoccurring CDR3 sequences is several orders of magnitudes lower. The negative control set is always the most diverse when comparing diversity between datasets, except for J family diversity, where the usage is slightly more uniformly spread across the data for the negative control TCRs than the epitope-specific TCRs. These results indicate a slightly restricted diversity of the epitope-specific TCR datasets when compared to the negative control set (figure 1).

**Fig. 1.**
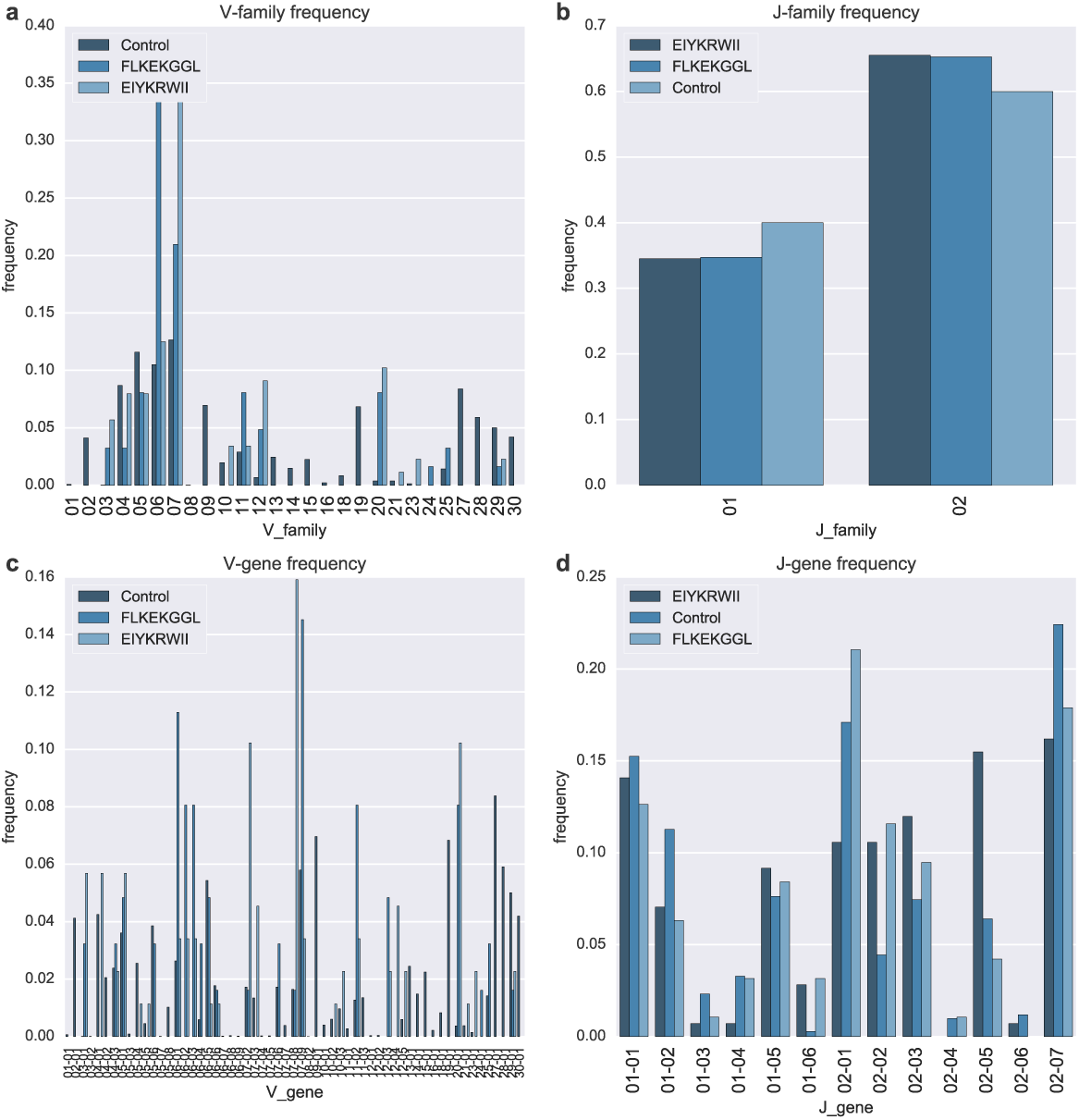
Segment usage across datasets. Segment usage is compared in terms of occurrence frequency at (a) V family, (b) J family, (c) V gene and (d) J gene level. Overall, no single segment distributions of segment usage are similar across different datasets, with slightly more restricted usage of segments in the peptide datasets, as is also reflected by the Shannon diversity (table 1)

**Table 1.** Descriptive statistics of all datasets. Diversity statistics were calculated based on the Shannon diversity.

| Peptide | EIYKRWII | FLKEKGGL | Control |
| --- | --- | --- | --- |
| total number of TCRBs | 142 | 95 | 56 |
| unique CDR3 sequences | 119 | 70 | 54 |
| CDR3 diversity | 6,82 | 5,92 | 15,71 |
| unique V families | 19 | 18 | 27 |
| unique V genes | 33 | 28 | 63 |
| unique J families | 2 | 2 | 2 |
| unique J genes | 12 | 12 | 14 |
| V gene diversity | 4,41 | 4,44 | 5,07 |
| V family diversity | 3,65 | 3,61 | 4,19 |
| J gene diversity | 3,17 | 3,18 | 3,16 |
| J family diversity | 0,93 | 0,93 | 0,97 |

### A highly performant classifier to distinguish two target epitopes

To test whether it is feasible to predict binding between a CD8+ T-cell’s TCRβ and an epitope, we first tested a ‘one-versus-one’, random forest classifier scheme where the classifier attempts to predict which of the two HIV epitopes a TCR sequence is most likely to be bound in a mutually exclusive way. The input features were derived from the V and J genes as well as the CDR3 sequence of the β chain of the TCR. The performance of this classifier was evaluated within a repeated subsampling validation approach in which part of the data is used as an independent test set for the trained classifier. This validation showed that the average classifier had a mean accuracy of 75.90% ± 5.45%, a mean AUC of 0.84 ± 0.05 and a mean PR of 0.81 ± 0.06 (FLKEKGGL) and 0.89 ± 0.04 (EIYKRWII) on independent test data. The high accuracy indicates that, in general, there is a high rate of both true positives and true negatives; in essence, the classifier is able to correctly assign to which peptide a given TCR will bind. As AUC values range from 1 (perfect prediction) to 0 (completely wrong prediction), with a value of 0.5 representing completely randomly assigned labels, the resulting average AUC value demonstrates that the classifier performs significantly better than random (one sample t-test; p < 0.001) (figure 2a). Finally, the mean PR, ranging between 0 (no true positives among predicted positives) and 1 (only true positives among predicted positives), demonstrates that the averaged classifier is able to retain a high predictive quality even under increasing numbers of predicted positives (figure 2b, 2c).

**Fig. 2.**
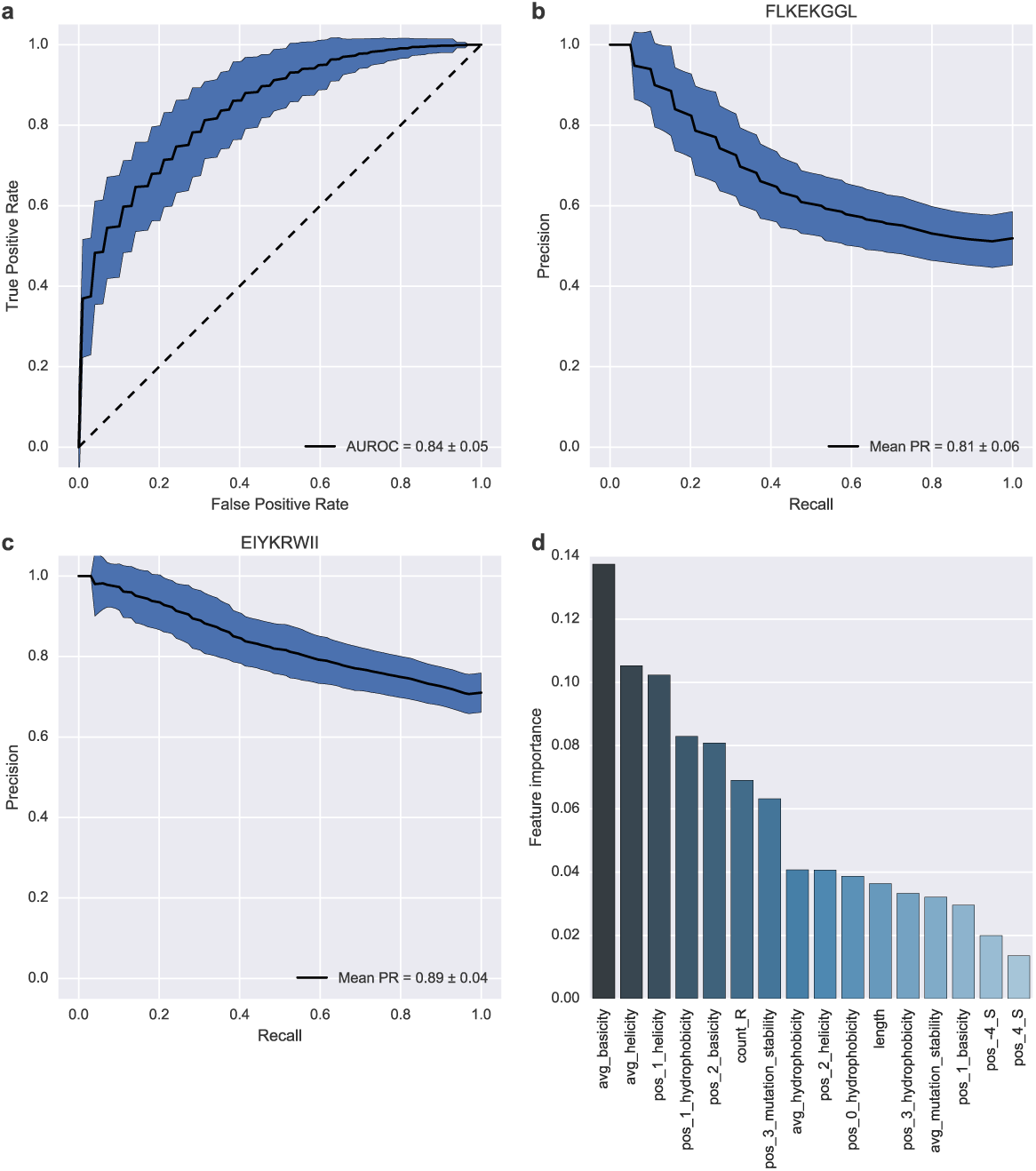
(a) Receiver-operating-characteristics curve or true positive rate versus false positive rate. Averaged values were plotted as a single line while the surrounding area indicates the standard deviation as observed during cross-validation. The striped diagonal indicates the performance of a random classifier where no distinction between the two peptides can be made based on TCR features. The more the average AUC curve is shifted towards the top left corner of the figure and away from the diagonal, the higher the performance of the evaluated classifier. (b, c) Precision versus recall. Averaged values were plotted as a single line while the surrounding area indicates the standard deviation as observed during cross-validation. In the perfect case, the PR curve is a horizontal line with precision always equal to 1, representing the lack of false positive predictions over the entire recall range. (d) Ranked feature importance. Features were generated based on the following TCR sequence derived characteristics: V gene, J gene, averaged CDR3 amino acid counts, positional CDR3 amino acid presence, averaged CDR3 amino acid physicochemical properties, and positional CDR3 amino acid physicochemical properties. Features are ranked on the x-axis according to their importance within the decision tree scheme of the random forest classifier as plotted on the y-axis

Features with high discriminatory power within the classifier can be supposed to play prominent roles within the biological recognition process between peptide and TCR. We thus investigated which features were most important during the model’s classification process. Despite the high number of features included in the classifier due to the positional encoding employed, only a limited number of features were assigned a large importance within the classification scheme (figure 2d). High-scoring features included averaged physicochemical properties (basicity, helicity, hydrophobicity) as well as physicochemical properties of the amino acids located slightly upwards of the centre of the CDR3 region (position 1 and 2). The only feature directly linked to a single amino acid is the overall arginine count within the CDR3. A notable absence of features encoding for single amino acids or V/J genes was observed among highly discriminatory features, even though they make up the bulk of the generated features.

### Epitope-specific TCRs can be picked out of a large TCR background

Evaluation of the ‘one-versus-one’ classifier scheme shows that differentiating between two peptides based on TCRβ sequence derived features is a feasible task. However, the scope of such a classifier remains limited in its applicability. To explore to which extend TCRβ sequence information can support sequence based TCR epitope predictors, we generalized the problem to identifying TCRβs that bind a given peptide from a larger set of TCRβs. This ‘one-versus-many’ scheme was applied and tested for both the FLKEKGGL and EIYKRWII peptide using non-epitope specific HLA-B*08-restricted, CD8+ TCR sequences as a negative control. While it is not known whether any of these control T-cell receptor sequences are capable of recognizing either B*08- FLKEKGGL or B*08-EIYKRWII, the upper limit of the expected abundance of T-cells that recognize a specific HLA-peptide combination has been estimated at 100 cells per million naïve T-cells (Jenkins and Moon 2012). As such, we assume that very few to none of these TCRβs are capable of interacting in a functionally relevant way with either of two HIV epitopes.

On the FLKEKGGL as well as the EIYKRWII peptide, the same pipeline was applied. TCRβ sequence features were generated in the same way as in the ‘one-versus-one’ classifier scheme. The performance of a classifier trained following the ‘one-versus-many’ scheme was evaluated within a repeated subsampling validation approach. Evaluation of the classifier revealed a mean accuracy of 93.78% ± 0.66%, a mean AUC of 0.80 ± 0.05 and a mean PR of 0.52 ± 0.06 for the EIYKRWII peptide. For the FLKEKGGL peptide, a mean accuracy of 94.45% ± 0.72%, a mean AUC 0.82 ± 0.05 of and a mean PR 0.61 ± 0.07 of were obtained. Evaluation metrics again indicate that both classifiers perform significantly better than random based on the AUC value (p < 0.001 for both the EIYKRWII and FLKEKGGL peptide) (figure 3 a, 3b) and reach similar performance levels as the ‘one-versus-one’ approach. The accuracy seemingly increases for the ‘one-versus-many’ classifiers as a consequence of class imbalance in the dataset (10 negative cases for every positive case) and is thus only marginally higher than the base accuracy of 0.91. PR values drop rapidly in comparison to the ‘one-versus-one’ scheme, likely also due to the class imbalance (figure 3c, 3d). With a higher number of negative classes, it becomes increasingly more difficult to retain a high predictive quality of the positive class and a low number of false positive predictions already severely affects precision. Given the increased complexity of the task, the slight drop in performance of the ‘one-versus-many’ classifiers is not completely unexpected. However, ‘one-versus-many’ classifiers are able to retain a high level performance level and reinforce the feasibility of creating more complex TCR epitope predictors. Differences in performance between the two ‘one-versus-many’ classifiers are likely to be a consequence of the number of patterns captured by the classifier that underlie recognition of the p-MHC by the TCR chain under investigation.

**Fig. 3.**
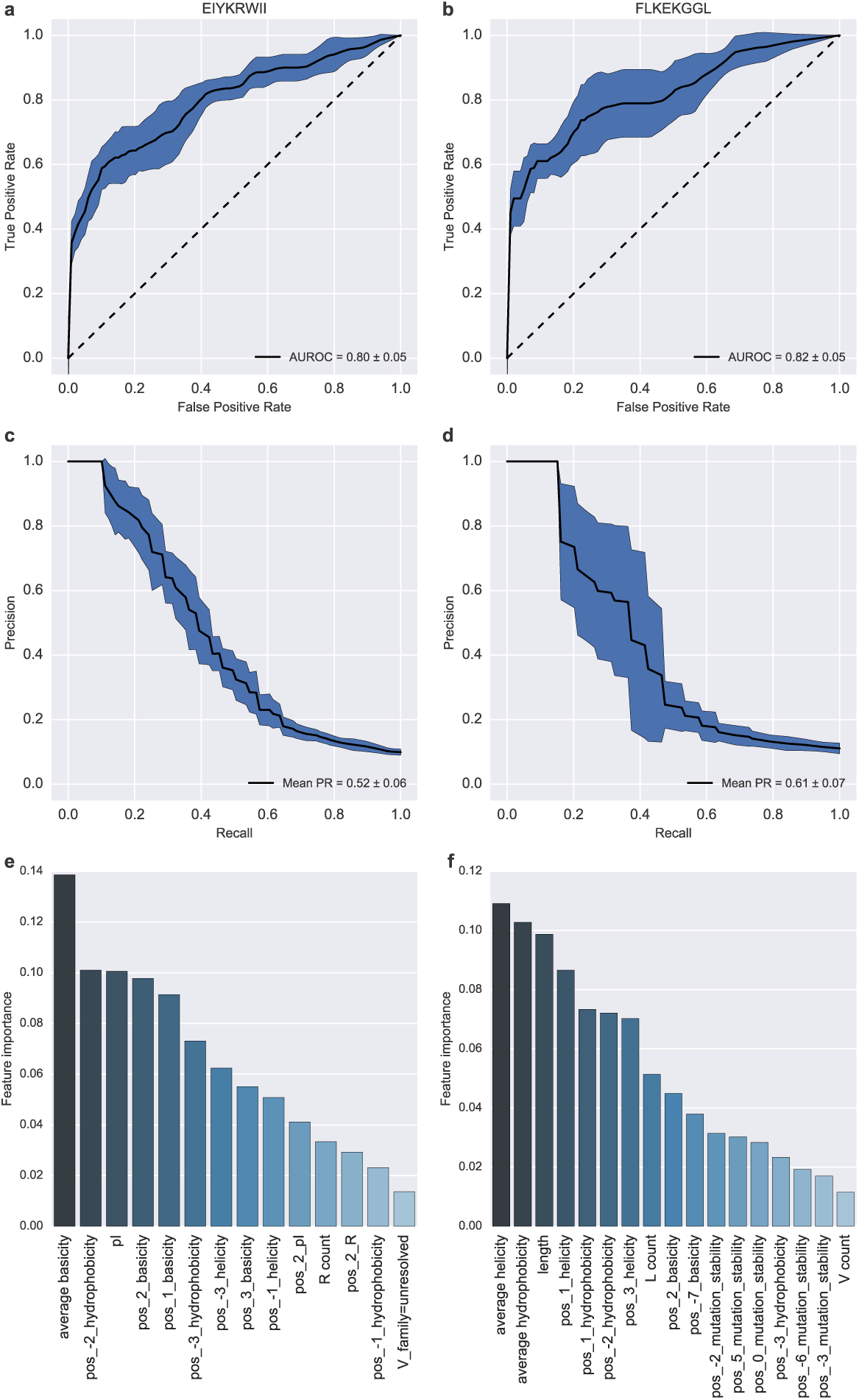
Comparison of the validation measures and feature importances for each peptide using the one-versus-many scheme. (a, b) Receiver-operating-characteristics curve or true positive rate versus false positive rate. Averaged values were plotted as a single line while the surrounding area indicates the standard deviation as observed during cross-validation. The striped diagonal indicates the performance of a random classifier where no distinction between the two peptides can be made based on TCR features. The more the average AUC curve is shifted towards the top left corner of the figure and away from the diagonal, the higher the performance of the evaluated classifier. (c, d) Precision versus recall. Averaged values were plotted as a single line while the surrounding area indicates the standard deviation as observed during cross-validation. In the perfect case, the PR curve is a horizontal line with precision always equal to 1, representing the lack of false positive predictions over the entire recall range. (e, f) Ranked feature importance. Features were generated based on the following TCR sequence derived characteristics: V gene, J gene, averaged CDR3 amino acid counts, positional CDR3 amino acid presence, averaged CDR3 amino acid physicochemical properties, and positional CDR3 amino acid physicochemical properties. Features are ranked on the x-axis according to their importance within the decision tree scheme of the random forest classifier as plotted on the y-axis

### Discriminatory features are similar in the one-versus-one or one-versus-many setting

To further explore the patterns captured by the classifier, highly discriminatory features were extracted from the classifier. Discriminatory features for the EIYKRWII peptide are focused on basicity and hydrophobicity, reflecting a similar pattern of features as found for the ‘one-versus-one’ scheme (figure 3e). Average basicity of the CDR3 amino acid sequence and basicity/hydrophobicity of amino acids near the centre of the CDR3 sequence seem to play the important roles next to the isoelectric point of the CDR3 sequence. The number of arginines within the CDR3 sequence was again found to be a discriminatory feature and potentially provides a more specific insight into the structural dimension of the interaction characteristics. While the FLKEKGGL peptide based classifier also relies heavily on hydrophobicity derived features of the CDR3 region to discriminate between binding and non-binding TCRs, helicity derived features seem to replace basicity derived features in importance. Similarly as to the most important features for the EIYKRWII peptide, these physicochemical properties seem to be concentrated near the centre of the CDR3 sequence (figure 3f). The CDR3 sequence’s length seems to be another important contributing factor together with the number of lysines in the CDR3 sequence. Overall, important features for both ‘one-versus-many’ based classifiers match well with those found for the ‘one-versus-one’ classifier, indicating that classifiers are able to faithfully retrieve important epitope-recognition patterns, even in different contexts.

### More TCR training samples result in a more performant classifier

Finally, to investigate the influence of the size of the training data on the performance both the ‘one-versus-one’ and ‘one-versus-many’ classifiers, models were trained with and without independent test data on increasing training data sizes. Regardless of the size of the training data, classifiers always performed with perfect accuracy if no independent test data was used, as can be expected from classification frameworks (figure 4, 5). In contrast, classifiers with independent test data benefited from increases in training data size and are likely to improve even further given a sufficiently large body of training data. As such, at least within the context of these classification schemes, the performance is likely to benefit from increases in experimental data.

**Fig 4.**
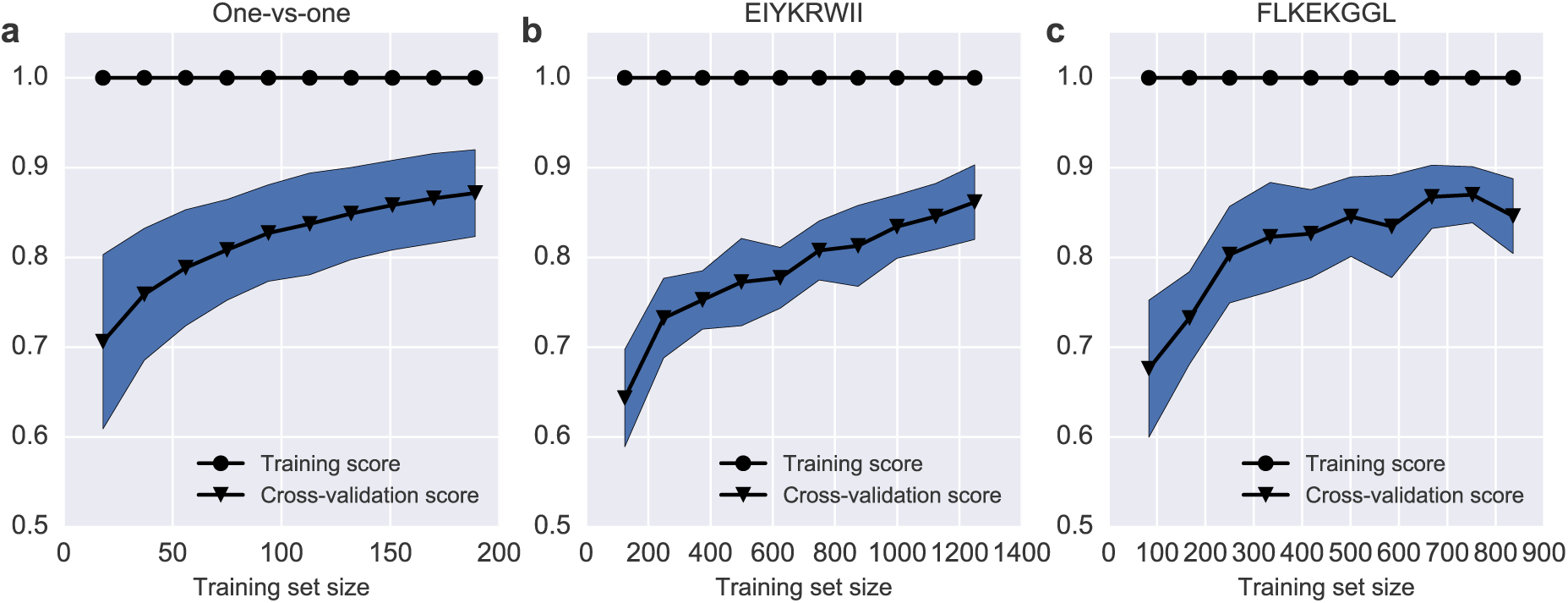
Learning curve for the different classifier schemes: (a) ‘one-versus-one’ scheme, (b) ‘one-versus-many’ scheme for the EIYKRWII peptide, (c) ‘one-versus-many’ scheme for the FLKEKGGL peptide. Influence of training data size on the accuracy of classifiers with access to all training data (blue dots) and cross-validated classifiers (green dots). AUC values were compared for 10 different training data sizes. Mean AUC values for fitted classifiers without independent test data are shown in blue while AUC values for cross-validated classifiers are shown in green. Surrounding areas of the curve indicate the variation in AUC values during cross-validation

## Discussion

In this paper, we set out to examine the feasibility of predicting epitope specificity from the sequence patterns contained within the TCR. Based on training data collected for 2 HIV-1 derived, B*08-restricted peptides, we trained random forest classifiers utilizing two different schemes to test whether TCR epitope prediction based on sequence level data is a feasible task. In the first, ‘one-versus-one’, scheme, the classifier was tasked to assign TCRs to either of two possible peptides. In the second, ‘one-versus-many’, scheme, classifiers were trained to find TCRs that bind to one specific peptide. In order to examine the properties that define the recognition of immunogenic peptides by a TCR, structural features were encoded representing the CDR3 amino acid sequence of the TCR β-chain as well as its respective V and J region.

As this is, to the best of our knowledge, the first attempt to tackle this problem at the TCR sequence level, it is not possible to compare its performance with other pre-existing classifiers. However, multiple performance evaluations indicate that the different classifiers performed reasonably well. The best performing scheme was the ‘one-versus-one’ scheme. However, the applications of this one-versus-one classifier are limited as it can only distinguish between the TCR sequences that bind one of two epitopes. Indeed, the importance of this classifier lies not in its immediate practical applicability, but rather in the framework it provides for future TCR-peptide recognition models and the insights that might be gained from these models. We can suppose that, given enough data for a large number of epitopes, this binary classifier can be expanded into a framework to predict the target epitope for any TCR sequence.

To demonstrate a more practical application, we incorporated negative control data from a large set of HLA-B*08-restricted TCRβs into ‘one-versus-many’ classification schemes. During these more difficult ‘one-versus-many’ classification schemes, performance was still high enough to prove that sequence level based models are capable of discriminating between epitope binders and non-binders. The good performance on held-out validation data, indicates that classifiers were likely able to capture real molecular features of the TCRβ CDR3 sequence that underpin the differential recognition of an epitope by a TCR within a B*08 context.

Of all the features generated, only a limited number of features had a high importance score within a given classifier. Supporting the likelihood that classifiers were able to consistently capture import features, high scoring features were generally shared across the different presented classifier setups. These features generally encoded either physicochemical properties averaged over the entire CDR3 region or physicochemical properties of amino acids located near its centre, with basicity and hydrophobicity as the most prominent physicochemical features. These findings correspond well with previous literature describing pMHC recognition by TCRs to be mediated by molecular interactions between the CDR loops and the pMHC complex, where the CDR3 loop plays a prominent role during epitope recognition (Jorgensen et al. 1992). As such, CDR3 loops with comparable physicochemical properties are likely to interact in similar ways with epitopes. In addition, the high rank of central physicochemical amino acid features suggests that these amino acids might be key in determining TCR specificity. The number of arginines in the CDR3 loop is one of two features capturing specific single amino acid data within the list of highly ranked features. Interestingly, arginine has previously been identified as a strongly conserved amino acid within the CDR3α loop of CD8+ T-cells recognising a HIV-1 epitope (Motozono et al. 2014).

Although the immediate applicability of the binary, ‘one-versus-one’ peptide classifier is limited in scope, its good performance illustrates the feasibility of creating high-performing p-TCR affinity models. We further demonstrate this feasibility by creating random forest classifiers that can distinguish TCRs that bind a specific epitope. Next to the binary classifier, these more general ‘one-versus-many’ classifier schemes set the stage for the development of more complex TCR prediction models in the future. Despite only using sequence information for the TCR β-chain, classifiers were able to differentiate their targeted epitope with high accuracy. In addition, the classifiers agree on highly discriminatory features, even for different classifiers contexts and are thus likely able to uncover important structural features. Thus it seems that the recognition determinants contained within the β-chain are already sufficient to predict epitope binding. These results do still leave room for increased performance. Indeed, learning curves for the different classifier schemes suggests that performance increases can still be gained by incorporating new training data. These results indicate that current models are therefore not necessarily bound by technical limitations but rather by a lack of suitable training data. As we anticipate the amount of available MHC-peptide-TCR data to increase in the future, we expect sequence based models to quickly gain in performance and become a valuable aid in future immunological studies. In particular, insights and advancements into TCR recognition of immunogenic epitopes might prove crucial in studies of auto-immunity, tumour susceptibility and vaccine design.

## Materials & Methods

### Data collection

Training data on T-cell receptor sequences and peptides were obtained from Costa et al. (2015). In this study, T-cells from chronically infected HIV-1 patients were stained with MHC tetramers and sorted by flow cytometry to select for CD8+ tetramer-positive T-cells. After extraction of mRNA from sorted T-cells, reverse transcription PCR was used to linearly amplify TCRβ chain sequences. PCR products were then transformed into *E. coli* bacteria, amplified and sequenced using capillary electrophoresis. From the data generated by Costa et al. (Costa et al. 2015), we collected TCRβ chain sequences from peptide-specific CD8+ T-cells for two HIV-1 derived HLA-B*08-restricted peptides (FLKEKGGL and EIYKRWII). In total, 95 TCRβ chains were collected for the FLKEKGGL peptide and 142 TCRβ chains for the EIYKRWII peptide.

Negative control data was obtained by querying the ImmuneACCESS database (https://clients.adaptivebiotech.com/immuneaccess) using the following terms: ‘human’, ‘TCRB’, ‘HLA-B*08’, ‘CD8+’ and ‘control’; which returned 66235 TCRβ chain sequences originating from a single individual. From these, 56023 unique, productive, in-frame sequences were withheld.

For each obtained TCRβ chain, the following information was collected: V family, J family, V gene, J gene, and CDR3 sequence; with V/J families and genes as defined by the Immunogenetics Information System (http://www.imgt.org) (Lefranc et al. 2015). D genes were not collected separately as the CDR3 sequence contains the D region’s sequence information.

### Feature creation

Several features were derived for each of the collected TCRβ chains. Each observed V or J gene was represented as a single feature. This feature was assigned a value of 0 or 1, representing respectively the absence or presence of the gene in a specific TCRβ chain. Because the V or J gene was not always available, the V and J family were also encoded in a similar way. The following properties were encoded as numeric features: the TCR CDR3 sequence length; the absolute count of each individual amino acid in the CDR3 sequence; the total mass of the amino acids in the CDR3 sequence; and the average CDR3 basicity, hydrophobicity, helicity, isoelectric point, and mutation rate. The average CDR3 mutation rate was calculated by taking the average of the mutation rate for each amino acid in the CDR3 sequence, where the mutation rate of an amino acid is obtained from the diagonal of a PAM250 substitution matrix. Physicochemical amino acid property values were used as described in MS2PIP (Degroeve et al. 2013). Positional features were added for each CDR3 residue position. Due to the variable length of the CDR3 sequences present in the data, amino acid positions were translated into numerical positions by assigning each position an index value relative to the centre of the CDR3 sequence. For example, a sequence of length 3 would be encoded by the positions -1, 0 and 1 whereas a sequence of length 4 would be encoded as -2, -1, 1 and 2. For each position encoding generated in such a way, a binary feature was created representing the presence or absence of an amino acid at that position was encoded in the same way as the V and J genes. In addition, numerical features encoding individual amino acid basicity, hydrophobicity, helicity, isoelectric point, and mutation stability were also created for each position. Features were always created prior to model training and evaluation.

### Model training & evaluation

Peptide binding was predicted using a random forest classifier (Breiman 2001) consisting of 200 trees, as implemented in Sci-kit learn (Pedregosa et al. 2011). To tackle the problem of predicting peptide binding to a TCR, a ‘one-versus-one’ scheme and a ‘one-versus-many’ scheme were employed. In the ‘one-versus-one’ scheme, the classifier was tasked with correctly assigning whether a TCR binds to either the EIYKRWII peptide or the FLKEKGGL peptide. In the ‘one-versus-many’ scheme, the classifier had to distinguish between TCRs that bind a given peptide and TCRs that don’t bind the given peptide. The ‘one-versus-many’ scheme was applied for both the EIYKRWII and the FLKEKGGL peptide.

During the ‘one-versus-one’ scheme, both the positive and negative class samples were considered to be of equal importance and their weight was set at 1. For the ‘one-versus-many’ scheme, class weights were set to be inversely proportional to the number of samples for that class to compensate for the larger amount of negative training samples. Other hyperparameters of the classifier were left at their default values, as random forests are highly performant classifiers that typically achieve excellent performance out of the box (Caruana et al. 2008). For the ‘one-versus-one’ scheme, the data was randomly subsampled 100 times into stratified 80%-20% training and testing data sets during model validation to provide a robust assessment of the classifier’s performance despite the limited amount of data available. In the ‘one-versus-many’ scheme, the classifier’s performance was assessed by creating 5 equally-sized, non-overlapping subsets from the positive data. For each positive subset, a subset of negative control samples equal to 10 times the amount of positive samples within the positive subset were randomly sampled without replacement from the negative control data. Positive and negative subset were then combined to generate a single fold. Training and test subsets were then created by using a single fold as test set while training the classifier on the remaining folds.

On each training set, feature selection was performed prior to training a new model using the Boruta algorithm (Kursa and Rudnicki 2010). For each new model, the following validation measures were calculated on the held-out test data: prediction accuracy, the area-under-the-receiver-operating-characteristic-curve (AUC), and the mean precision over a recall range of 0 to 1 (PR). Overall classifier performance was evaluated in terms of prediction accuracy, AUC values, and mean PR values averaged over 100 random stratified training-test sets for the ‘-one-versus-one’ scheme and over 10 folds for the ‘one-versus-many’ scheme. The receiver-operating-characteristic curve and precision-recall curve were drawn up for each model as well. Here, precision is interpreted as how many of the TCRs predicted to bind peptide 1 actually bind peptide 1 and recall as how many of the TCRs that bind peptide 1 are predicted to bind peptide 1. Because PR values are computed for the peptide with 1 as label, for the ‘one-versus-one’ scheme, the PR values were also calculated with reversed labels to obtain PR values for both peptides. The evaluation measures were reported as their mean over the number of subsampled executions ± their standard deviation.

Feature importance was evaluated based on the Gini importance (Hastie et al. 2009) to provide an overview of each feature’s ability to discriminate between the two peptides. In addition, classification accuracy was evaluated by drawing learning curves for increasing sizes of training data with and without independent test data. Classifiers with independent test data used a stratification and sampling scheme as described above for their respective schemes while classifiers without independent test data were tested on their training data.

All data and code used within this manuscript can be found in the following GitHub repository: https://github.com/bittremieux/TCR-classifier.

### Statistical analyses

Statistical analyses were performed in Python. The Shannon-Wiener diversity was calculated using the Sci-kit bio package (http://scikit-bio.org/). Statistical results were considered significant whenever the (corrected) p-value < 0.05.

## Acknowledgments

This research was funded by the University of Antwerp [BOF Concerted Research Action] and the Research Foundation Flanders (FWO) [Personal PhD grants to NDN (151316), PMo (1141217N), BC (11O1614N)]

## References

Breiman L (2001) Random forests. Mach Learn 45:5–32. doi: 10.1023/A:1010933404324

Calis JJA, Maybeno M, Greenbaum JA, et al (2013) Properties of MHC Class I Presented Peptides That Enhance Immunogenicity. PLoS Comput Biol 9:e1003266. doi: 10.1371/journal.pcbi.1003266

Caruana R, Karampatziakis N, Yessenalina A (2008) An empirical evaluation of supervised learning in high dimensions. Proc 25th Int Conf Mach Learn - ICML ’08 96–103. doi: 10.1145/1390156.1390169

Costa AI, Koning D, Ladell K, et al (2015) Complex T-Cell Receptor Repertoire Dynamics Underlie the CD8 T-Cell Response to HIV-1. J Virol 89:110–9. doi: 10.1128/JVI.01765-14

Degroeve S, Martens L, Jurisica I (2013) MS2PIP: A tool for MS/MS peak intensity prediction. Bioinformatics 29:3199–3203. doi: 10.1093/bioinformatics/btt544

Hastie T, Tibshirani R, Friedman J (2009) The Elements of Statistical Learning. Elements 1:337–387. doi: 10.1007/b94608

Jenkins MK, Moon JJ (2012) The role of naive T cell precursor frequency and recruitment in dictating immune response magnitude. J Immunol 188:4135–40. doi: 10.4049/jimmunol.1102661

Jensen PE (2007) Recent advances in antigen processing and presentation. Nat Immunol 8:1041–1048. doi: 10.1038/ni1516

Jorgensen JL, Esser U, Groth BF de S, et al (1992) Mapping T-cell receptor-peptide contacts by variant peptide immunization of single-chain transgenics. Nature 355:224–230. doi: 10.1038/355224a0

Krangel MS (2009) Mechanics of T cell receptor gene rearrangement. Curr. Opin. Immunol. 21:133–139.

Kursa MB, Rudnicki WR (2010) Feature Selection with the Boruta Package. J Stat Softw 36:1–13. doi: Vol. 36, Issue 11, Sep 2010

Lefranc MP, Giudicelli V, Duroux P, et al (2015) IMGT R, the international ImMunoGeneTics information system R 25 years on. Nucleic Acids Res 43:D413–D422. doi: 10.1093/nar/gku1056

Meysman P, Ogunjimi B, Naulaerts S, et al (2015) Varicella-Zoster Virus-Derived Major Histocompatibility Complex Class I-Restricted Peptide Affinity Is a Determining Factor in the HLA Risk Profile for the Development of Postherpetic Neuralgia. J Virol 89:962–969. doi: 10.1128/JVI.02500-14

Motozono C, Kuse N, Sun X, et al (2014) Molecular Basis of a Dominant T Cell Response to an HIV Reverse Transcriptase 8-mer Epitope Presented by the Protective Allele HLA-B*51:01. J Immunol 192:3428–34. doi: 10.4049/jimmunol.1302667

Mustafa AS (2013) In silico analysis and experimental validation of mycobacterium tuberculosis-specific proteins and peptides of mycobacterium tuberculosis for immunological diagnosis and vaccine development. Med. Princ. Pract. 22:43–51.

Pedregosa F, Varoquaux G, Gramfort A, et al (2011) Scikit-learn: Machine Learning in Python. J Mach Learn Res 12:2825–2830.

Rossjohn J, Gras S, Miles JJ, et al (2015) T cell antigen receptor recognition of antigen-presenxting molecules. Annu Rev Immunol 33:169–200. doi: 10.1146/annurev-immunol-032414-112334

Soria-Guerra RE, Nieto-Gomez R, Govea-Alonso DO, Rosales-Mendoza S (2015) An overview of bioinformatics tools for epitope prediction: Implications on vaccine development. J. Biomed. Inform. 53:405–414.

Stranzl T, Larsen MV, Lundegaard C, Nielsen M (2010) NetCTLpan: Pan-specific MHC class I pathway epitope predictions. Immunogenetics 62:357–368. doi: 10.1007/s00251-010-0441-4

Turner SJ, Doherty PC, McCluskey J, Rossjohn J (2006) Structural determinants of T-cell receptor bias in immunity. Nat Rev Immunol 6:883–894. doi: 10.1038/nri1977

Vita R, Overton JA, Greenbaum JA, et al (2015) The immune epitope database (IEDB) 3.0. Nucleic Acids Res 43:D405–D412. doi: 10.1093/nar/gku938

